# The fly liquid-food electroshock assay (FLEA) reveals opposite roles for neuropeptide F in avoidance of bitterness and shock

**DOI:** 10.1101/2020.02.10.942508

**Authors:** Puskar Mishra, Shany E. Yang, Austin B. Montgomery, Addison R. Reed, Aylin R. Rodan, Adrian Rothenfluh

**Author notes:** **Author contributions:** P.M. and A.R. designed research; P.M. performed experiments and designed electrical boards, data acquisition and analysis tools; S.E.Y. and A.R.Re. performed experiments; A.B.M. worked on software tools; A.R.Ro. and A.R. procured funding; P.M., S.E.Y., A.B.M., A.R.Ro., and A.R. wrote the paper. The authors declare no competing interests.

## Abstract

Proper regulation of feeding is important for an organism’s well-being and survival. Food intake in *Drosophila* can be determined in a number of ways, including by measuring the time a fly’s proboscis interacts with a food source in the fly liquid-food interaction counter (FLIC). Here, we show that electrical current flowing through flies during this interaction is aversive and leads to a reduction in food intake. Based on the FLIC, we engineer a novel assay, the fly liquid-food electroshock assay (FLEA), which allows for current adjustments for each feeding well. Using the FLEA, we show that both external incentives as well as internal motivational state can serve as drivers for flies to overcome higher current (electric shock) to obtain superior food. Unlike similar assays in which bitterness is the aversive stimulus for the fly to overcome, we show that current perception is not discounted as flies become more food-deprived. The FLEA is therefore a novel assay to accurately measure incentive motivation in *Drosophila.* Using the FLEA, we also show that neuropeptide F is required for proper perception or processing of an electroshock, a novel function for this neuropeptide involved in processing of external and internal stimuli.

**Significance Statement:** Many neuropsychiatric disorders, such as depression or addiction, are associated with alterations in motivated behavior. Assays measuring incentive motivation determine how driven an organism is to attain a goal, like food, or how attractive an incentive is. These tests require the animal to put effort into obtaining the reward, which can include physical work or overcoming an aversive stimulus. Such assays for *Drosophila* feeding have relied on flies overcoming bitterness to obtain their food. However, the perception of bitterness is discounted as flies become food deprived, confounding the interpretation. Here, we developed a novel assay that does not suffer from the same shortcomings and thus allows for more accurate assessments of incentive motivation in this widely used model organism.

## INTRODUCTION

Motivation can be regarded as an organism’s goal-directed quest for change, for example the search for food in the face of starvation. Numerous human conditions show aberrations in motivated behaviors, such as psychiatric disorders like depression or addiction, but also neurodegenerative diseases such as Parkinson’s or dementia (1). The mechanistic dissection of the neural and molecular mechanisms of motivated behaviors is thus of considerable relevance to human health. Assays to measure motivation in animal models generally involve them exerting physical effort, or overcoming an aversive stimulus, such as walking across an electrified grid. When the external incentive is increased, rats for example, will cross a grid that delivers a larger electric shock (2). In addition to external incentives acting as motivators, the other main component to motivational behavior is the internal state and resulting drive of the animal (2). The integration and valuation of internal drive and external incentive is what stirs the animal into goal-directed action. Feeding is one of the fundamental actions in animals and is normally under tight regulation to keep an organism’s energy expenditure and stores in balance. Eating disorders are common human dysregulations of feeding and are still not well understood at the molecular and neural level. Since all animals regulate their food intake, model organisms can help in the dissection of the mechanisms regulating the motivation of feeding behavior.

The vinegar fly, *Drosophila melanogaster*, has been a genetic model organism for over 100 years, and numerous assays exist to determine a fly’s feeding behavior. These include measuring food consumption from a tiny capillary (3), or lacing the food with quantifiable substances, such as dyes (4, 5), radioactive compounds (6), or even oligonucleotides (7). Recently, additional assays have been developed that rely on feeding flies closing an electrical circuit that allows the interaction between fly and food to be measured in intensity and duration (8). Even though the latter assays do not measure actual ingestion, the time spent interacting with the food correlates well with the amount ingested (9). Thus, these assays are valuable additions due to of their wide temporal range – from milliseconds to days – over which they can record feeding events. Some of the above feeding assays have been coupled with bitter substances, in order to determine flies’ willingness to overcome aversion to get to food, resulting in a measure of their feeding motivation (10). However, as flies become starved, their peripheral perception of bitterness decreases (11, 12). Thus, seemingly increased motivation can be caused, at least in part, by the decreased perception of the aversive stimulus in the first place, thereby confounding the quantitative assessment of flies’ motivation.

Here, we develop a novel feeding assay based on the FLIC (fly liquid-food interaction counter; (8)). Our novel assay allows for individual feeding wells to be paired with different amounts of current delivered, enabling us to ask what variables motivate flies to overcome a higher current to obtain food. We show that external incentives and internal drive both act as feeding motivators. Lastly, we find that neuropeptide F (npF) also plays a role, albeit in the perception of the electrical current itself, thus revealing a novel function for this neuropeptide.

## RESULTS

The fly liquid-food interaction counter (FLIC) is a *Drosophila* feeding assay that allows for continuous online feeding monitoring (8). When flies standing on a metal plate make contact with the liquid food, they complete an electrical circuit, which allows for precise measurement of the duration the flies interact with the food. In addition, the amplitude of the signal depends on whether flies touch the food with their legs (lower amplitude “leg events”) or engage in food consumption using their proboscis (higher amplitude “proboscis events”).

We wanted to determine whether the FLIC can be used to measure feeding-bout duration under various conditions. When we determined the median duration of feeding-bouts in a 15-min FLIC assay, we found that the duration increased with the length of prior food deprivation (Fig. 1 *A*). Similarly, feeding-bout duration also increased when we increased the amount of sucrose offered (Fig. 1 *B*, left). After an 18-hr food deprivation, the median feeding-bout length was 3.2 sec on 400 mM sucrose. We similarly food deprived flies for 18 hr and then filmed them feeding on liquid sucrose in a petri dish. When we determined the duration of the first feeding bout, we again saw that bout duration increased with the amount of sucrose offered (Fig. 1 *C*). We also found that the median first bout lasted 12 sec on 64 mM sucrose, considerably longer than in the FLIC on the higher, 400 mM, sucrose concentration.

**Fig. 1.**
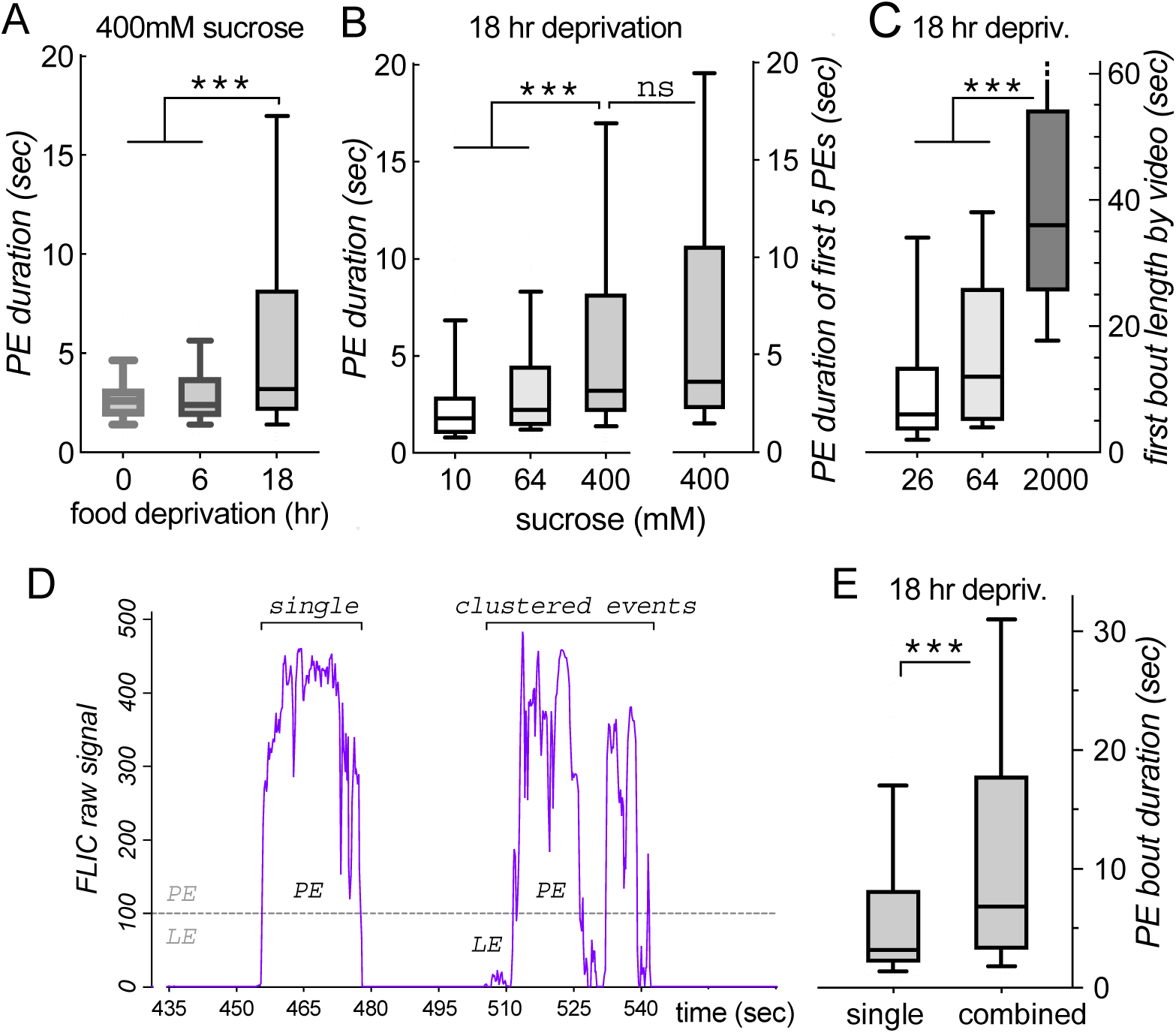
Feeding bout length measured in the FLIC. *(A)* Longer food deprivation times result in significantly longer proboscis events (PE) on 400 mM sucrose (\*\*\**p* < 0.001, Mann-Whitney test with Dunn’s correction, n = 233, 233, 349 events. Here, and in following panels, the first 15 minutes of FLIC events were analyzed, unless specified otherwise). *(B)* PE durations are longer when flies are offered higher sucrose concentrations (left side; \*\*\**p* < 0.0001 n = 75, 137, 349). If only the first 5 PEs per feeding well are analyzed (right side), the median PE duration increases slightly, but not significantly (ns = not significant, *p* = 0.37, n = 349, 143). *(C)* Length of first feeding bout as measured by video recording. The median length of bouts, here defined as uninterrupted engagement with the liquid food, increase with sucrose concentration offered (\*\*\**p* < 0.0001, n = 45, 58, 46). Note that the bouts are considerably longer when compared to measurements in the FLIC *(A,B). (D)* Example data from a FLIC well (400 mM sucrose, 18 hr deprivation). Some PEs occur in single isolation (left). Others come in clustered bouts of PEs (right), together with leg events (LE, signal amplitude < 100 over baseline), that follow in close succession. *(E)* When PEs that occur in groups of events (with an inter-event interval of less than 5 sec) are grouped into combined feeding events, the median bout length increases significantly compared to analyzing all PEs as single bouts (as done in *A*; *p* < 0.0001, n = 349, 188; all data are shown as medians with quartile boxes and 10-90 percentile whiskers).

To try to understand this discrepancy in bout duration, we first tested the hypothesis that later bouts in the 15-min FLIC assay were shorter, as flies became satiated, thus lowering the measured median bout length. Analyzing only the first 5 bouts in the FLIC — from a total of 10 flies — revealed a small, but not significant increase in bout duration (Fig.1 *B*, right). We thus rejected shortness of later bouts as the cause for the bout duration difference in the FLIC versus feeding in a dish. In our free-feeding filming experiment, we counted a brief disengagement of the proboscis, followed by immediate re-engagement of the proboscis with the food, as being part of one and the same feeding bout, reasoning that there was no interruption of the feeding by a distinctly different behavior. In the FLIC, proboscis interaction-bouts sometimes appear as long and isolated, and sometimes in clusters, interspersed with leg interactions (Fig. 1 *D*). When we examined the frequency distribution of the bout duration, we saw an obvious inflection point at 5 sec (*SI Appendix*, Fig. S1). We therefore grouped interaction-bouts containing at least one proboscis interaction that were closer than 5 sec into one long bout. This led to a significant increase in the median bout duration in the FLIC on 400 mM sucrose, from 3.2 to 6.8 sec (Fig. 1 *E*). Considering bout structure and grouping is therefore part of the reason why the bout duration in the FLIC is shorter than when free feeding.

However, even the bout-grouped median duration of 6.8 sec on 400 mM sucrose in the FLIC (Fig. 1 *E*) was still considerably shorter than the free-feeding median of 12 sec on 64 mM sucrose (Fig. 1 *C*). We therefore tested a third hypothesis, which posited that the current in the FLIC is causing a reduction in proboscis-interaction. To test this, we added a fluorescent dye to the sucrose solution in the FLIC and then measured the amount ingested in a plate reader, as we have done before (5). Indeed, turning on the FLIC current caused a significant reduction in food intake of both 64 and 400 mM sucrose (Fig. 2 *A*). This confirmed that the FLIC current is aversive to flies when they are feeding.

**Fig. 2.**
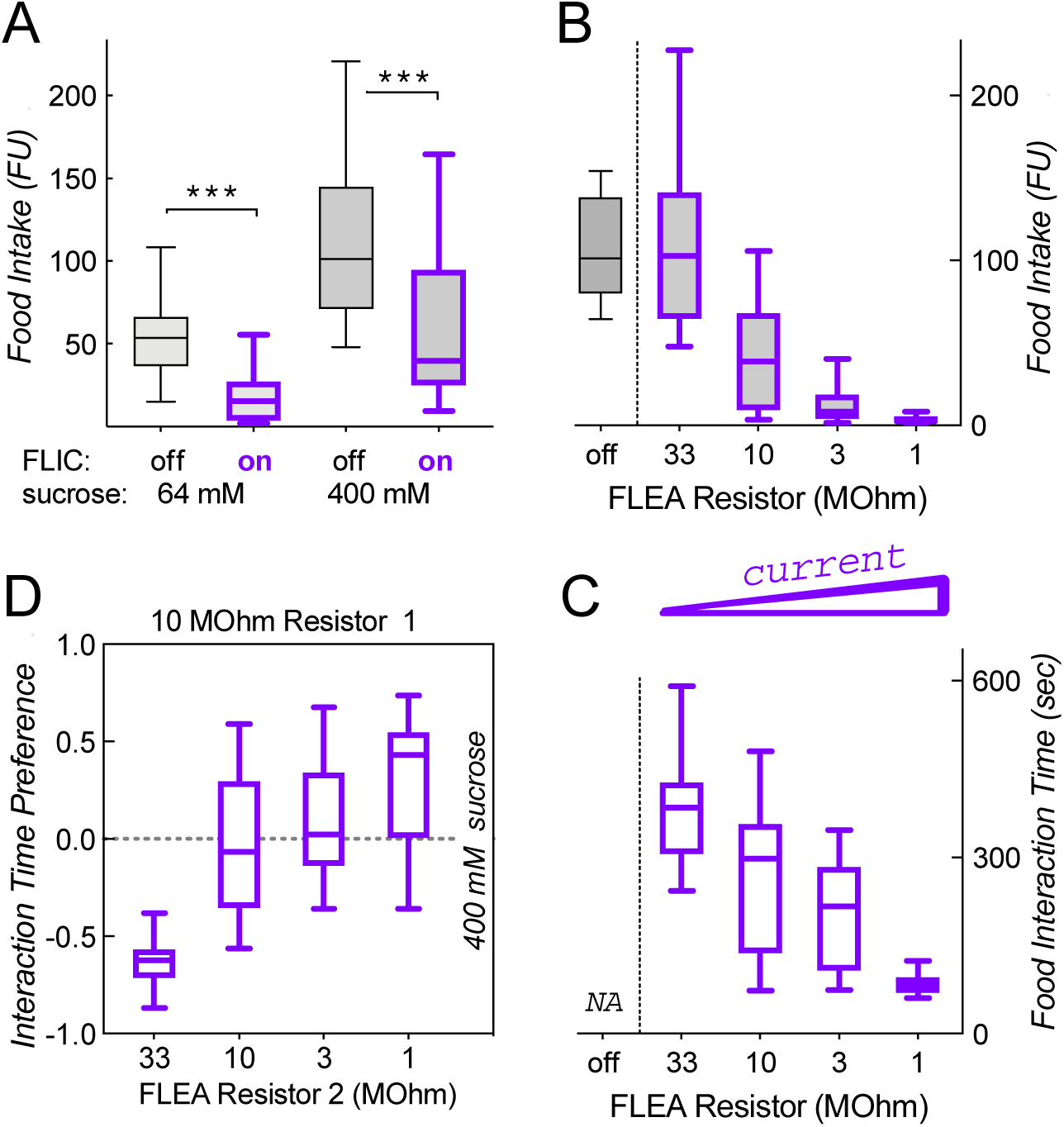
FLIC current is aversive to flies. *(A)* Turning on FLIC current significantly reduces food intake of 18-hr food deprived flies, as measured by fluorescent dye ingestion (FU = fluorescence units measured in plate reader; \*\*\**p* < 0.0001, n = 33–51). *(B)* Redesigned FLIC with adjustable current (FLEA) shows a decrease in food intake as a function of current (*p* < 0.0001; one-way Kruskal-Wallis ANOVA, n = 28–39). Note that the original FLIC contains a 10 MOhm resistor. *(C)* Food interaction time, as measured by the FLEA current, is also significantly reduced as the current increases (*p* < 0.0001; n = 11–12; NA = not available). *(D)* Flies change their interaction time preference as the current increases in one of two equal-sucrose wells (*p* < 0.0001; n = 10–11).

The FLIC has a fixed design, which includes a 10 MOhm resistor to limit the current flow when the circuit is closed (8). We redesigned the FLIC in a way that allowed us to modularly exchange this current-limiting resistor for each food well. Because this new assay also allowed us to increase the current, we named it the FLEA, for fly liquid-food electroshock assay. We first tested whether altering current flow when flies closed the circuit would have an impact on sucrose intake labeled with fluorescent dye. As hypothesized, the smaller the resistor, and the higher the current, the lower the amount of food ingested (Fig. 2 *B*). Similarly, the time spent interacting with the food as measured by the current signal in the FLEA was also shorter, the higher the current (Fig. 2 *C*).

We reasoned that we might make use of the current as an aversive stimulus and designed the FLEA as a 2-choice assay where one choice goes with higher current. We tested whether flies would prefer to interact with food that was paired with the lesser current, while food quality remained equal. Indeed, the flies’ interaction preference changed as we altered the current-limiting resistor in one of the two wells (Fig. 2*D*), suggesting that we might be able to use the FLEA as an assay to measure feeding motivation. Such feeding experiments—asking whether flies are willing to overcome an aversive stimulus—have been described using bitter substances mixed in with one of the two feeding solutions (10). The willingness to overcome bitterness can then serve as a proxy for flies’ feeding motivation. However, an assay including bitterness has a significant confounder: the perception of bitterness depends on the flies’ food deprivation status, with hungry flies showing less bitter acuity (11, 12). We confirmed this by testing flies’ willingness to overcome 1 µM denatonium and indeed found significantly reduced aversion to this bitter substance with longer periods of food deprivation. Flies demonstrated less avoidance (less negative interaction time preference) after 18 hr compared with shorter deprivation (Fig. 3 *A*). We also determined the preference for proboscis (feeding) and leg (tasting) interactions separately, and both showed an effect with increased food deprivation (Fig. 3 *B*). This makes sense, since there are bitter sensory neurons located on both the legs and the proboscis (12, 13). Our data thus confirmed that bitterness is discounted by food deprivation. To use the FLEA as an assay for feeding motivation, the perception of the current should not be altered by the duration of prior food deprivation. We therefore performed the same experiment as with denatonium, this time with current as the deterrent, varying the duration of prior food deprivation. Avoidance of the well with higher current increased with deprivation time (Fig. 3 *C*). This would suggest that flies actually become more sensitive to current as they are food deprived for longer. However, when we analyzed the proboscis and leg interaction preference separately, neither of them depended on the duration of food deprivation (Fig. 3 *D*). Flies strongly avoided proboscis interaction with the higher current, while leg interactions were insensitive. This suggested two things: first, in a setting of 10 vs 33 MOhm (Fig. 3 *D*), the leg-mediated current cannot be perceived, probably because it is considerably smaller than proboscis-mediated current (see Fig. 1 *D*). Second, because food deprivation increases the frequency of proboscis over leg events (as flies are hungry and want to feed), proboscis interactions become more prevalent after 18 hr of food deprivation. As the proboscis interactions are more sensitive to current than the leg interactions, this skews the total (proboscis+leg) interaction preference towards the negative, i.e lower-current well, explaining the apparent increase in sensitivity to current of total event preference with increasing food deprivation (Fig. 3 *C*).

**Fig. 3.**
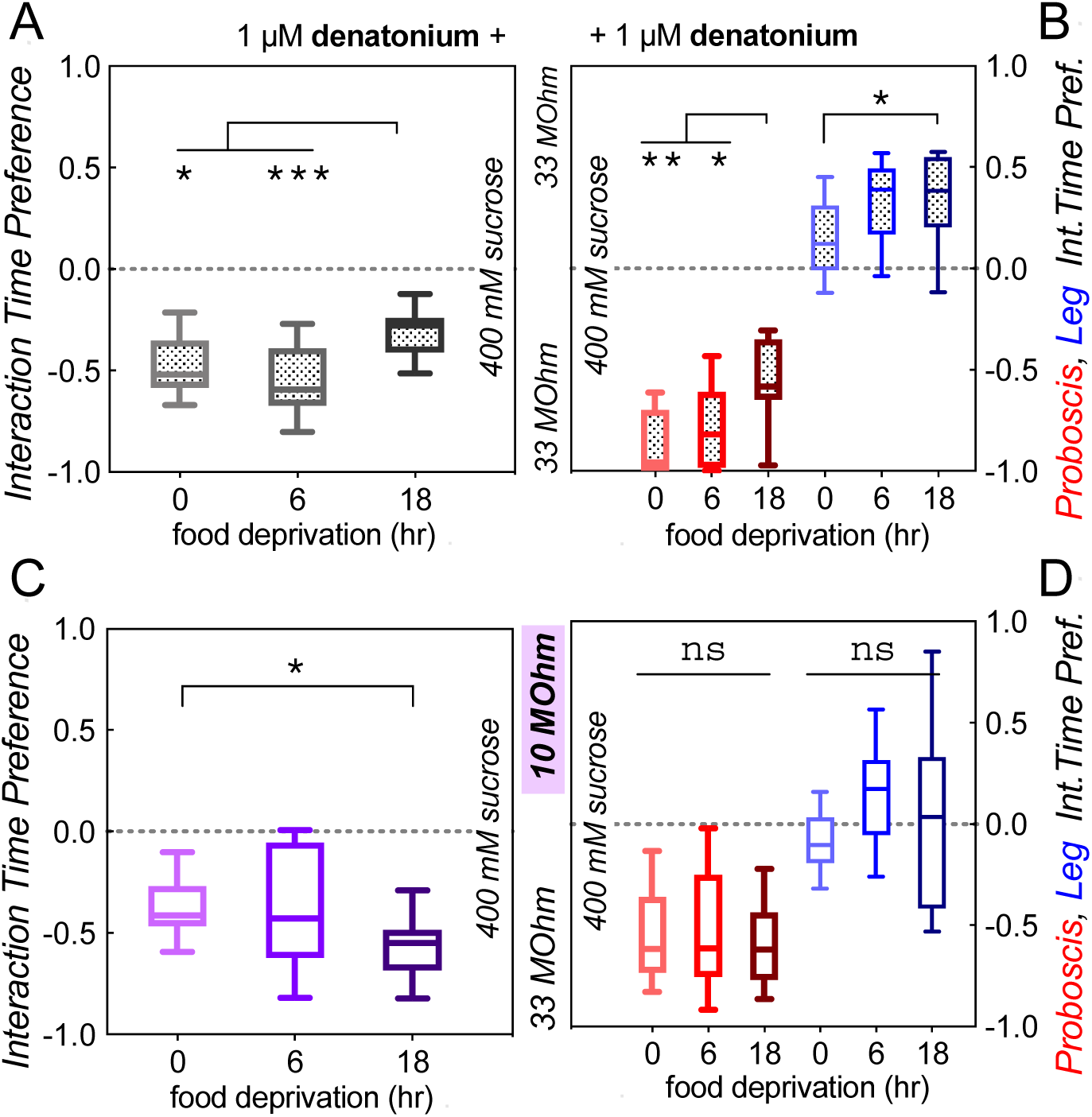
Unlike bitterness, current is not discounted upon food deprivation. *(A)* 18-hr deprived flies show reduced avoidance of denatonium (\*\*\**p* = 0.0011, \**p* = 0.0495, Kruskal-Wallis test with Dunn’s correction). *(B)* Both leg (blue, \**p* = 0.031) and proboscis (red, \*\**p* = 0.0013, \**p* =0.039, n = 13–15) interactions show decreased sensitivity to denatonium with increased food deprivation. *(C)* Current avoidance increases with food deprivation when determining total (proboscis+leg) interaction time (\**p* = 0.022). *(D)* However, neither proboscis (red, ns = not significant, *p* = 0.73), nor leg (blue, ns *p* = 0.06, n = 15–20) interaction time preference changes with food deprivation. The apparent decrease in total interaction preference in *(C)* is caused by a shift from leg to proboscis interactions upon food deprivation, lowering the combined Preference Index.

Because our data suggested that current perception by the proboscis and by the legs is not discounted by food deprivation, we wanted to establish the FLEA as an assay for motivation, and we next tested whether an external incentive would induce flies to overcome a higher current. As hypothesized, 18 hr food-deprived flies showed less aversion to a higher current when the high-current well contained more sucrose (Fig. 4 *A* and *B*). Thus, flies are willing to overcome current, if enough of an external incentive is paired with it. Furthermore, the FLEA allowed us to assign a value on the incentive, which is the incentive size attractive enough to equal the aversion to a given current, resulting in a preference index of 0. In this experiment, it took a four-fold increase in sucrose concentration to offset the aversion to 10 MOhm current (Fig. 4).

**Fig. 4.**
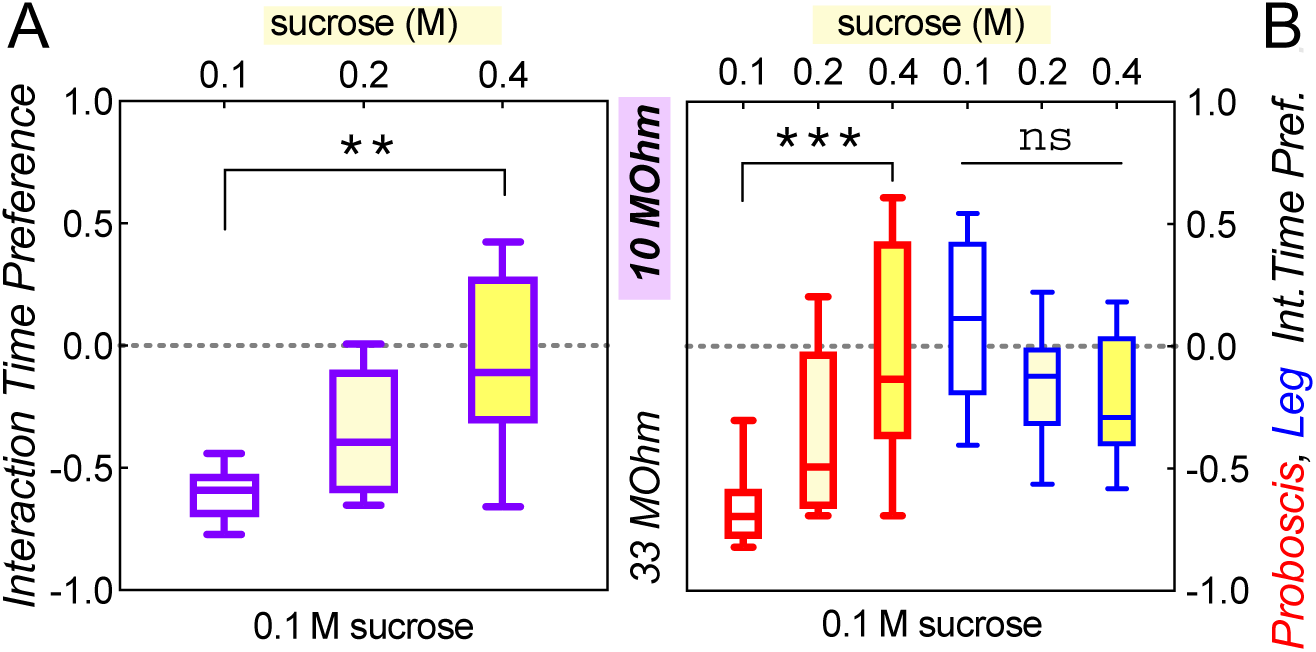
An external sucrose incentive causes reduced current avoidance in food-deprived flies. *(A,B)* Increasing the sucrose concentration in the food well with higher current reduces flies’ avoidance of that food well. Both total interaction preference (*A*, \*\**p* = 0.002, Kruskal-Wallis test with Dunn’s correction), as well as proboscis interaction preference (*B*, \*\*\**p* = 0.0009, n = 10–12) increase significantly with sucrose concentration. Flies were food-deprived for 18 hours before testing.

Next, we wanted to test whether internal drive would induce flies to overcome higher current. To do so, we compared flies that were food-deprived for 6 vs. 18 hr, a time difference that has been shown to lead to significant behavioral changes (14). When we first performed this experiment pairing 10 mM sucrose with 33 MOhm current versus 100 mM sucrose with the 10 MOhm current, we found a slight trend, but no significant effect of food deprivation (data not shown). Using the 33 MOhm resistor leads to a current that does not deter flies from feeding (Fig. 2 *B* and *C*), thereby setting up a steep gradient against 10 MOhm current. We decided to instead use a resistor pair where both currents were perceptible to the flies. In a choice of these currents, each paired with 100 mM sucrose, flies preferred to interact with food paired with the lower 20 MOhm over the higher 4.7 MOhm current (Fig. 5 *A*). The median preference index in this setting was -0.19 (Fig. 5 A), which was considerably less aversive compared to our prior 33 MOhm vs. 10 MOhm comparisons with equal sucrose, where the preference indices ranged from -0.41 to -0.62 (Fig. 2 *D*, 3 *C*, 4 *A*). This confirmed that our 20 vs. 4.7 MOhm setup presented a lesser current gradient then the initial 33 vs. 10 MOhm choice. As before (Fig. 3 *C* and *D*), the perception of the current in and of itself did not depend on the 6 vs. 18 hr duration of food deprivation (Fig. 5 *A* and *B*). When we next paired 100 mM sucrose with the higher 4.7 MOhm current, this solution was equally palatable to 6-hr deprived flies as a 10 mM/20 MOhm pairing. However, after an additional 12 hr of food deprivation, the flies preferred the 100 mM/4.7 MOhm well (Fig. 5 *C* and *D*), suggesting that an increased internal feeding drive caused the flies to be willing to overcome a higher current to obtain better food.

**Fig. 5.**
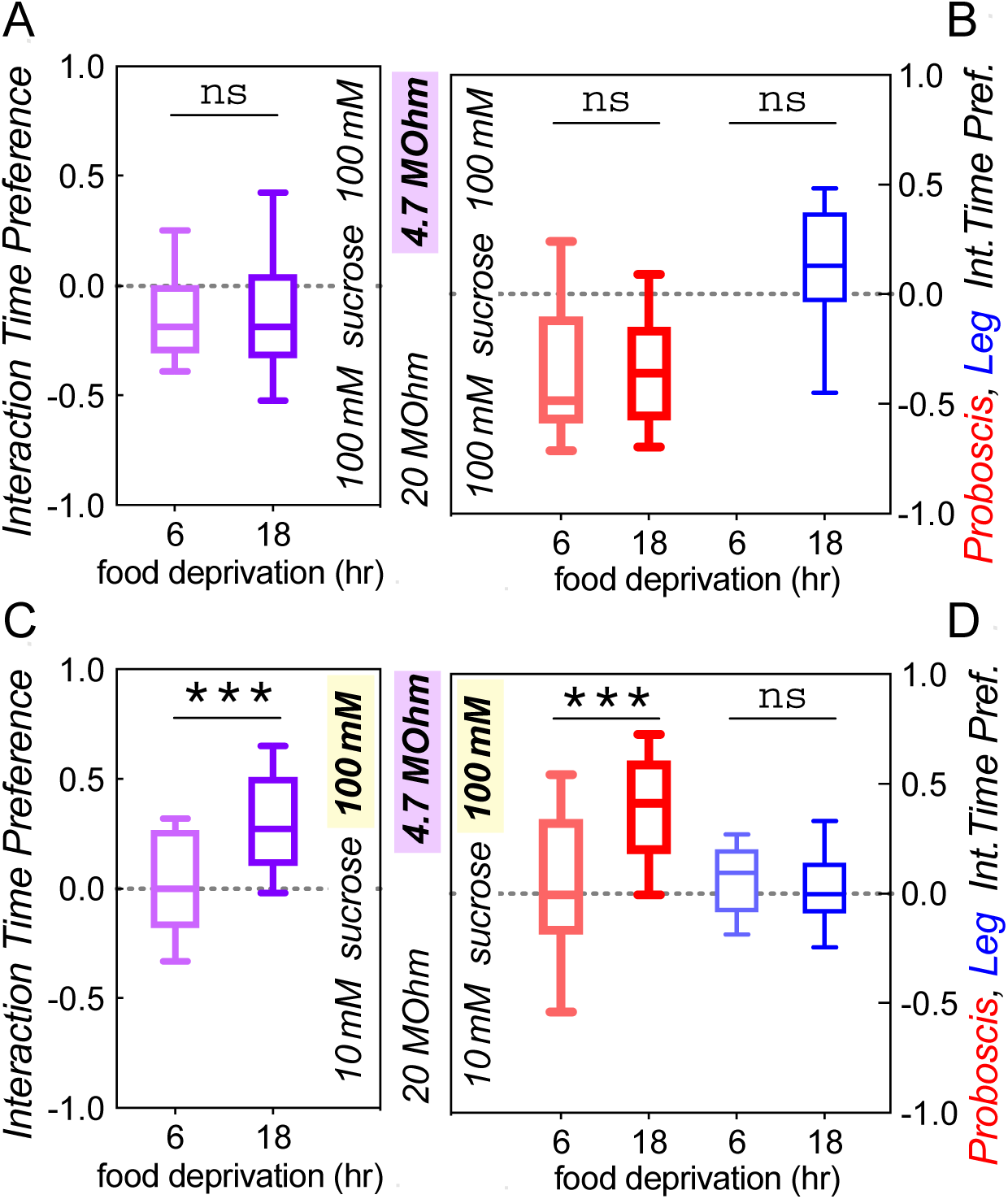
Increased internal feeding drive causes reduced current avoidance. *(A)* Increasing the duration of food deprivation from 6 to 18 hr has no effect on flies’ avoidance of 4.7 vs. 20 MOhm current at equal sucrose (ns *p* = 0.88, Mann-Whitney U test). *(B)* This was also true for proboscis and leg interaction preference (ns *p* = 0.50 and 0.42, n = 27, 28). (C) The combination of 100 mM sucrose with 4.7 MOhm current became attractive only after 18 hr of food deprivation (\*\*\**p* = 0.0005). (D) This was also evident in the proboscis interaction preference (\*\*\**p* = 0.0005), while leg interaction preference remained unchanged (ns *p* = 0.38, n = 26, 30).

Lastly, we wanted to test the role of the neuropeptide F (npF) in feeding motivation, using our novel FLEA assay. Fly larvae showed an enhanced willingness to ingest bitter food with increased npF signaling, while reduced npF signaling made larvae ingest less bitter-laced food (15). This suggested that npF is involved in feeding motivation and that npF signaling might similarly cause flies to overcome higher current to get to better food. To our surprise, when we silenced npF neurons by overexpression of the inwardly rectifying Kir2.1 channel in *npF-Gal4* neurons, those flies were more attracted to the higher current side (10 MOhm/100 mM sucrose vs. 33 MOhm/10 mM; Fig. 6 *A*), the opposite result of what we expected. We then tested whether these flies were as sensitive to the current itself as their controls, and we found that they were less deterred by current when presented with equal sucrose in both wells (100 mM, 10 vs. 33 MOhm; Fig. 6 *B*). This suggested that npF is required for proper perception of electroshock, a function for npF not previously proposed. We therefore wanted to replicate this finding using *npF-Gal4* driving a temperature sensitive *shibire*^*ts*^ gene causing neuronal silencing. Larvae carrying *npF*>*shi*^*ts*^ were previously shown to be more sensitive to quinine in the food at the restrictive temperature. We first wanted to replicate this finding in adult flies. Using our two-choice fluorescence consumption assay (5), we found that at the control temperature, 18 hr food-deprived *npF*>*shi*^*ts*^ flies preferred 100 mM sucrose/7 mM caffeine vs. 50 mM sucrose alone. However, at the restrictive 32° temperature these flies avoided the sucrose/caffeine solution (Fig. 6 *C*), consistent with the proposed model that npF is required to overcome bitterness in food (15). We then tested these flies in the FLEA, and again found that reduced npF signaling at the restrictive temperature lowered flies’ avoidance to higher current at equal sucrose (Fig. 6 *D*). This again supported the hypothesis that npF signaling is involved in the perception of electroshock. Next, we assessed proboscis bout duration with varying current. There was no effect of silencing npF neurons (*npF*>*shi*^*ts*^ flies) when current is imperceptible (33 MOhm resistor). However, at higher currents (10 mOhm resistor), *npF*>*shi*^*ts*^ flies showed a significantly increased median proboscis-bout duration at the restrictive temperature (Fig. 6 *E*). This suggested that npF signaling is required to inform flies of the aversive shock, which induces termination of a feeding bout. We also replicated this finding using npF receptor mutants, *npfR*^*c01896*^, which also showed significantly longer proboscis-bout duration when exposed to greater current (10 MOhm resistor), but not on low/imperceptible current (33 MOhm; Fig. 6 *F*). Therefore, three distinct genetic *npF* manipulations supported the interpretation that npF signaling is required for proper perception of electroshock.

**Fig. 6.**
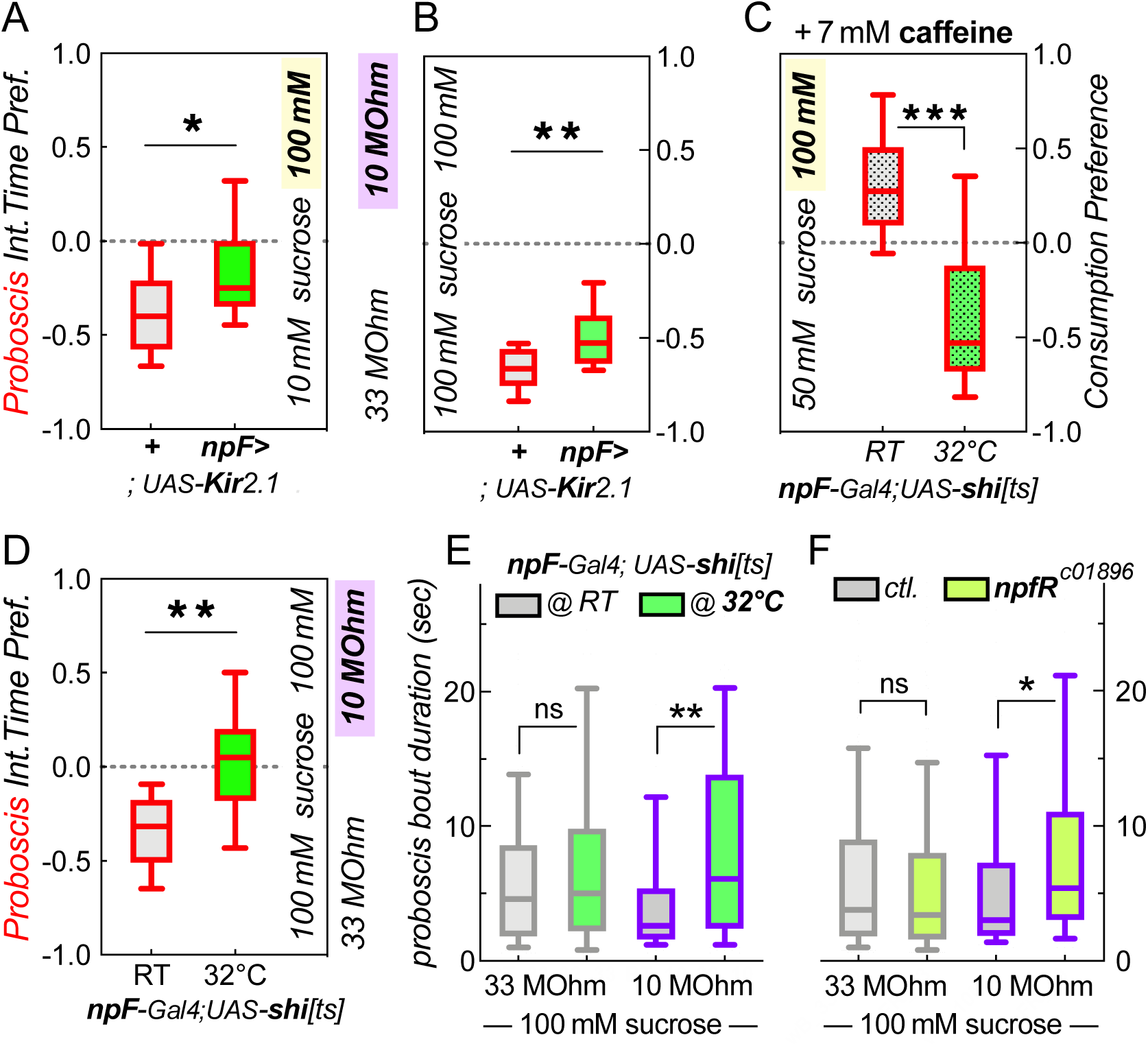
Reduced npF siganling leads to reduced shock-avoidance. *(A)* Flies with reduced npF signaling show decreased avoidance of a high current/high sucrose combination (\**p* = 0.017, Mann Whitney U-test, n = 18). *(B)* These flies also show reduced avoidance of higher current at equal sucrose concentration combination (\*\**p* = 0.006, n = 13–14). *(C)* 18 hr food-deprived flies with reduced npF signaling show increased avoidance of a bitter caffeine/high sucrose combination at the restrictive temperature in a fluorescence ingestion choice assay combination (\*\*\**p* < 0.0001, n = 15–16). (D) These same flies show decreased avoidance of a higher current combination (\*\**p* < 0.003, n = 9–11). *(E)* Reduced npF signaling also leads to an increase in median proboscis bout length at the restrictive temperature with 10 MOhm current (\*\**p* = 0.0012, n = 47, 86), but not on the 33 MOhm, imperceptible current well (ns *p* = 0.14, n = 103, 77). *(F)* Similary, mutation in the npF receptor leads to increased bout duration on the 10 MOhm (** *p* = 0.007, n = 96, 80), but not 33 MOhm well (ns *p* = 0.20, n = 255, 271).

## DISCUSSION

Here, we describe the FLEA as a novel feeding assay based on the design for the FLIC, where flies touch a liquid food source and complete an electrical circuit, leading to a small current (8). This allows for the precise measurement of feeding-time interactions and can be used longitudinally, over the course of days (8). We were interested in more short-term measurements to determine the variables affecting individual feeding bouts. As expected, feeding bouts were lengthened with increasing food quality, and prior food deprivation (Fig. 1 *A* and *B*). Because the absolute durations of these feeding bouts were considerably smaller than what we observed by filming freely feeding flies (Fig. 1 *C*), we suspected that the FLIC current might actually be aversive to the flies. Indeed, the FLIC current limited by a 10 MOhm resistor caused about a three-fold reduction in actual food ingestion (Fig. 2 *A*). Our data also showed that a 33 MOhm current was undetectable by the flies (Fig. 2 *B*), and they preferred to interact with a 33 MOhm feeding well over a 10 MOhm well, at over a 3:1 ratio (Fig. 2 *D*, 3 *C*, 4 *A*). The negative value of a 10 MOhm current is only offset by a four-fold increase in sucrose concentration. The current generated from a leg interaction is about 5 times smaller than that for a proboscis interaction (Fig. 1 *D*), and our data suggest that flies cannot detect the 10 MOhm current with their legs (Fig. 3 *D* and 4 *B*). This difference is of similar magnitude as the 10 MOhm aversive / 33 MOhm imperceptible current ratio, and overall our data suggest that while the 10 MOhm FLIC current is close to innocuous, it is aversive to flies touching the food with their proboscis and will report skewed durations of feeding interactions. Higher currents lead to even shorter food interactions and smaller volumes ingested (Fig. 2 *B* and *C*), but even at the highest, and very aversive 1 MOhm current, we found no evidence of flies being electrocuted.

Based on these findings, we re-engineered a FLIC-like assay that allows for adjustable currents to be paired with each one of two feeding wells in order to measure flies’ feeding motivation. Prior assays paired one well with a bitter substance, to gauge flies’ motivation to overcome an aversive stimulus while feeding (10). One confound in this setup is that flies’ devalue bitterness with increasing food deprivation (Fig. 3 *A* and *B*; (11)). This makes sense in the wild, where a (hungry) “beggar can’t be a chooser”, but it also means that assays of feeding motivation relying on bitterness as a deterrent are confounded by the flies’ internal state of satiety. Thus, a fly’s willingness to overcome a bitter substance is a combination of its internal deprivation state, or drive, plus a peripheral reduction in the perception of the bitterness in the first place (12). For the FLEA to be an improved measure of incentive motivation, we needed to show that the perception of the current would not change as a function of the internal feeding drive. Indeed, we found that increasing food deprivation from 6 to 18 hr did not alter flies’ avoidance of higher current (Fig. 3 *D* and 5 *B*), while it did reduce their avoidance of bitter denatonium (Fig, 3 *A* and *B*). Using the FLEA, we then found that both increasing an external incentive (higher sucrose concentration, Fig. 4), as well as increasing flies’ internal drive (longer food deprivation, Fig. 5 *C* and *D*) would induce them to overcome a larger current to obtain a higher quality food source. The FLEA therefore represents an improved *Drosophila* assay that can be used to quantitate incentive motivation in this highly manipulable model organism.

The npF neuropeptide has previously been shown to be important for larvae to overcome bitter-laced food (15). We replicated these results in adult flies, where we found that reduced npF signaling made flies less willing to overcome bitter caffeine to obtain a preferable sucrose solution (Fig. 6 *C*). However, when we performed the equivalent experiment with high vs. low sucrose/current pairings, loss of npF had the opposite effect and made flies more willing to overcome higher current (Fig. 6 *A*). Control experiments with 3 distinct npF manipulations revealed that decreased npF signaling reduced flies’ perception of current (Fig. 6). Our findings do not invalidate previous findings indicating that npF is involved in overcoming aversive stimuli in order to get superior food. However, we were unable to assess this, as we discovered here that npF is required for normal perception of the electroshock. The FLEA therefore revealed a novel function for this neuropeptide. Other than npF’s involvement in feeding motivation and overcoming bitterness (15), npF is also involved in the regulation of feeding and sleep, where increased npF signaling leads to more feeding and reduced sleep (16). It also acts as a gate for the retrieval of appetitive memories, ensuring that flies remember these memories when they are hungry (17). Furthermore, flies prefer to spend time in a place where their npF neurons are optogenetically activated (18). All these results are consistent with the model that npF is largely involved in motivation for positively reinforcing behaviors. However, male flies that are sexually frustrated – by lack of mating and continued rejection from already mated females – prefer to drink more alcohol (19). In that paradigm, frustrated males drink even more if they lack npF signaling, and increased npF signaling reduces their alcohol preference compared to controls. Thus, npF seems to mediate alcohol aversion (5, 19), possibly by enhancing the sensation of aversive stimuli. An involvement for npF in stimulus sensation/processing is also suggested by experiments showing that the L1 npF neurons are involved in peripheral olfactory sensitivity to ethyl butyrate (20). There is therefore precedent for npF’s involvement in not just internal motivation, but also in processing of external stimuli. We here reinforce that idea, by using our novel fly liquid-food electroshock assay, to show that npF signaling is required for the proper perception/processing of an aversive external electroshock.

## METHODS

### Fly Husbandry and Behavior

Male flies, age 2-8 adult days were used for all experiments. Flies were grown and kept on standard cornmeal/agar medium at 25°C with 70% relative humidity. Male *w* Berlin* flies were used as controls. Transgenic flies were outcrossed to the *w* Berlin* genetic background for at least 5 generations. Food deprivation was done in vials containing 0.7% agar only, as a water source. Freely feeding flies were filmed in a petri dish with a liquid sucrose drop containing 0.3% blue #1 to ensure we only analyzed the first feeding bout when scoring the movies. The FLIC assays were performed as described (8). For the design of the FLEA, see SI Methods, but in brief, each of 8 feeding wells in the FLEA contains a modular slot for a small board with distinct size current-limiting resistors, which can be placed independently of each other. FLEA data was acquired with from arenas with 2 wells per arena.

### Data Analysis and Statistics

The FLIC data was analyzed as described (8), in brief, signal of amplitude >100 was designated a proboscis event, and smaller amplitude interactions were deemed leg events. We also filmed flies in the FLIC and a correlation of the filmed behavior with the obtained FLIC signal and designations suggested a sensitivity and specificity of ∼90% for distinguishing leg from proboscis events. In the FLEA, the signal amplitude is changed as a function of the current limiting resistor, and we normalized the data accordingly, such that we obtained the same 0–1023 data range. The *Interaction Time Preference Index* was calculated by taking the difference in total time interacted between the two wells and dividing it by the sum of the time interacted with both wells. This yields an index of 0, if both wells are equally interacted with, and +1 or –1 if only one well was interacted with exclusively. For detailed processing and analysis of the FLIC and FLEA signal and data, see SI Methods. All data were checked for normality using Prism 8 (GraphPad Software Inc., San Diego, CA). Data were not normally distributed, and we excluded outliers for data sets with n > 8 if they fell 1.5x the interquartile range outside of the upper and lower quartiles. Data were compared using Mann Whitney U-tests, for pairwise comparisons, and Kruskal-Wallis tests with Dunn’s correction for multiple comparisons.

## ACKNOWLEDGEMENTS

We thank lab members for critical discussion of experiments and the manuscript. Flies were obtained from the Bloomington stock center. This work was funded by the NIH R01DK110358 (A.R.Ro.), NIH R01AA019526, R01AA026818 (A.R.), U2M2 (A.R.Ro., A.R.) and the University of Utah Neuroscience Initiative (A.R.).

## Supplementary Information

Supplementary Methods

Figure S1

### SUPPLEMENTARY METHODS

#### FLEA system overview

The FLEA system is comprised of five components: feeding unit, resistor modules, data acquisition device, NI LabVIEW software, and analysis software in R. The first component, the feeding monitoring unit is composed of a conductive metal baseplate, plastic reservoir, and printed circuit board. It houses eight feeding wells to conduct behavioral experiments. The second component, the resistor modules, is responsible for supplying current to each well in the feeding monitoring unit. The resistor modules are customizable to each of the 8 feeding wells and can be modularly exchanged. The third component, NI USB 6001 DAQ, is responsible for detecting analog signals from eight wells and forwarding the signal to the fourth component, NI LabVIEW software. The LabVIEW data acquisition software, NI SignalExpress, allows modification and customization of all the parameters of the system and record the data. For the last component, the R statistical analysis software is used for analyzing and visualizing the data and time preference analysis for behavioral experiments.

#### FLEA hardware

The behavior board are composed of an aluminum plate, a plastic food reservoir, and a plastic cover based on the design of the FLIC. A custom-made circuit board (AutoDesk EAGLE design available upon request) allows for the use of 8 wells and has receiving slots for 4 current resistor modules. The printed circuit board, the resistor modules, and the NI USB 6001 DAQ contain all the electronics needed for signal recognition, power supply to the board, and signal forwarding contacts to a computer for data acquisition. The printed circuit board contains two non-inverting operational amplifiers to provide a systemic gain of approximately 1.2, and it also contains 2×6 board-to-board male contacts for the attachment of resistor modules to the printed circuit board. The resistor module is the component that supplies current each well. It contains two customizable current limiting resistors, one for each well of a 2-well choice arena, and four gain resistors: two per well. The current limiting resistors are used to regulate the amount of electrical current permitted to pass through the flies.

#### FLEA signal processing and analysis

FLEA raw signal data is sampled at 500 Hz. A simple low-pass filter was utilized with the window size of 100 to reduce noise. Then, one mean data point was generated from the filtered signal for every 100 points to reduce the sampling rate from 500 Hz to 5 Hz. The process of filtering and sample rate reduction results in better resolution than sampling at 5 Hz. Then, filtered data was converted to the same scale as FLIC readings intensities ranging from 0 to 1023. The conversion factor for each current-limiting resistor was determined by measuring signal amplitude from circuits closed by defined resistors, standing in for flies. Baseline intensity varies linearly and/or nonlinearly through time. For a linear baseline, the baseline estimation was performed by computationally estimating zero-slope baseline. For non-linear baseline, we implemented a non-linear, non-parametric baseline adjustment algorithm – local polynomial regression (Loess; (21)) – to computationally estimate the baseline. Loess is a locally weighted polynomial regression performed via iterations of an M-estimation procedure with tricube kernel and Tukey’s biweight function as weighting parameters for time and intensity (21). Window span and polynomial degree of regression were specified independently for each dataset. Baseline correction of FLEA data was then performed by subtraction of the estimated baseline. Residuals of baseline estimation were removed by zeroing values less than 4au intensity.

Each detected peak was classified as a leg event (LE, maximal intensity <100) or proboscis event (PE, maximal intensity ≥100). Additionally, we identified exclusion criteria to eliminate false positive events (like food splatter), device errors, and to better correlate feeding events with flies’ observed behavior. Exclusion criteria for number of events per assay were derived from large pooled datasets from various conditions with the cutoffs based on the mean ± 2.5 standard deviations (eg. between 4 and 212 events for a 30-min assay). Similarity, the exclusion criteria for event duration was calculated based on the third quartile plus 2x the inter-quartile range (which came to 4 and 40 seconds, for leg and proboscis events, respectively).

**Supplementay Figure 1.**
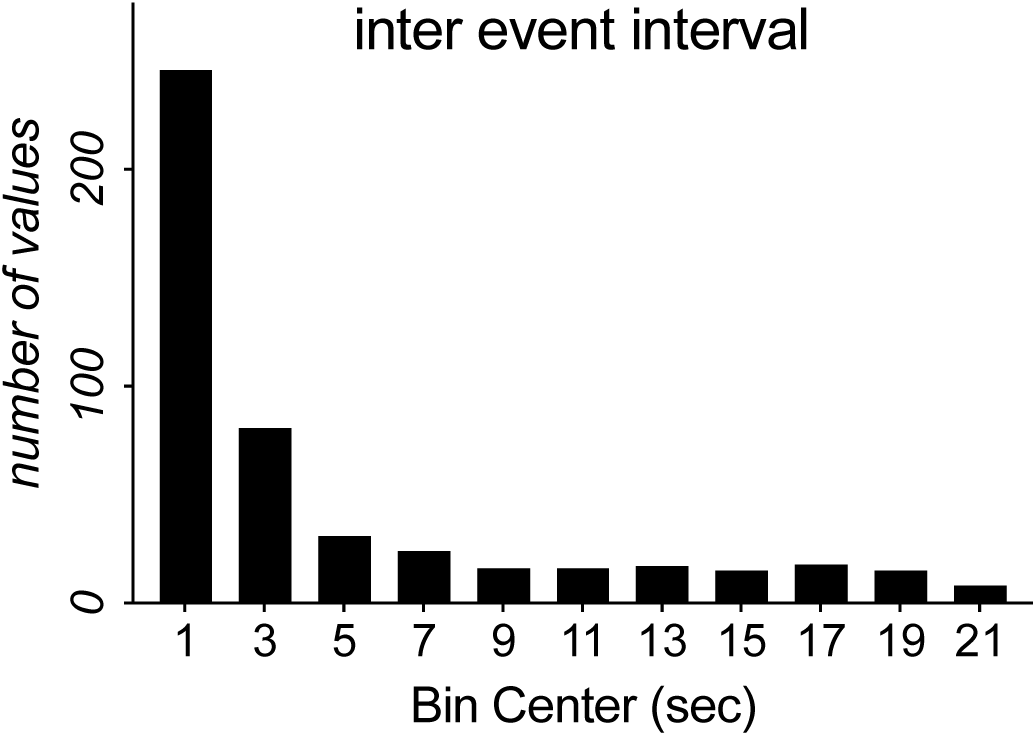
Frequency plot of inter-event intervals from the FLIC (related to Fig. 1 *D*). We chose 5 sec, the inflection point of this distribution, to group events together, or apart.

